# Some remarks on moments for stochastic chemical kinetics

**DOI:** 10.1101/021410

**Authors:** Eduardo D. Sontag, Abhyudai Singh

## Abstract

We analyze a class of chemical reaction networks for which all moments can be computed by finite-dimensional linear differential equations. This class allows second and higher order reactions, but only under special assumptions on structure and/or conservation laws.

## 1 Preliminaries on stochastic chemical kinetics

We start by reviewing standard concepts regarding master equations for biochemical networks, see for instance [3].

Chemical systems are inherently stochastic, as reactions depend on random (thermal) motion. Deterministic models represent an aggregate behavior of the system. They are accurate in much of classical chemistry, where the numbers of molecules are usually expressed in multiples of Avogadro’s number, which is *≈* 6 × 10^23^. In such cases, basically by the law of large numbers, the mean behavior is a good description of the system. The main advantage of deterministic models is that they are comparatively easier to study than probabilistic ones. However, they may be inadequate when the “copy numbers” of species, i.e. the numbers of units (ions, atoms, molecules, individuals) are very small, as is often the case in molecular biology when looking at single cells: copy numbers are small for genes (usually one or a few copies), mRNA’s (in the tens), ribosomes and RNA polymerases (up to hundreds) and certain proteins may be at low abundances as well. Analogous situations arise in other areas, such as the modeling of epidemics (where the “species” are individuals in various classes), if populations are small. This motivates the study of stochastic models. We assume that temperature and volume Ω are constant, and the system is well-mixed.

We consider a chemical reaction network consisting of *m* reactions which involve the *n* species

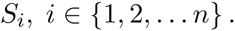

The reactions *R*_*j*_, *j ∈ {*1, 2, *…, m}* are specified by combinations of reactants and products:

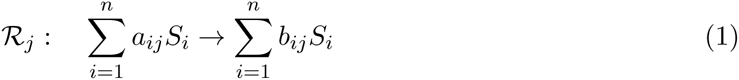

where the *a*_*ij*_ and *b*_*ij*_ are non-negative integers, the *stoichiometry coefficients*, and the sums are understood informally, indicating combinations of elements.

The data in (1) serves to specify the stoichiometry of the network. The *n × m stoichiometry matrix* Γ = *{γ_ij_}* has entries:

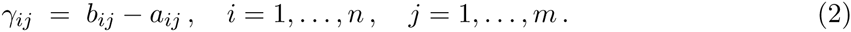

Thus, *γ*_*ij*_ counts the net change in the number of units of species *S*_*i*_ each time that reaction *R*_*j*_ takes place. We will denote by *γ*_*j*_ the *j*th column of Γ:

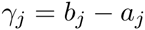

where

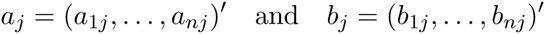

(prime indicates transpose) and assume that no *γ*_*j*_ = 0 (that is, every reaction changes at least one species).

In general, for every 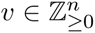, we denote *⊕v* = *⊕*(*v*_1_, *…, v_n_*) := *v*_1_ + *…*+ *v*_*n*_. In particular, for each *j ∈ {*1, *…, m}*, we define the *order* of reaction *R*_*j*_ as

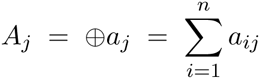

(the total number of units of all species participating in the reaction). One allows the possibility of zero order, that is, for some reactions *j*, *a*_*ij*_ = 0 for all *i*. This is the case when there is “birth” of species out of the blue, or more precisely, a species is created by what biologists call a “constitutive” process, such as the production of an mRNA molecule by a gene that is always active. Zeroth order reactions may also be used to represent inflows to a system from its environment. Similarly, also allowed is the possibility that, for some reactions *j*, *b*_*ij*_ = 0 for all *i*. This is the case for reactions that involve degradation, dilution, decay, or outflows.

Stoichiometry information is not sufficient, by itself, to completely characterize the behavior of the network: one must also specify the *rates* at which the various reactions take place. This can be done by specifying “propensity” or “intensity” functions.

### 1.1 Stochastic models of chemical reactions

Stochastic models of chemical reaction networks are described by a column-vector Markov stochastic process *X* = (*X*_1_, *…, X_n_*)^*′*^ which is indexed by time *t ≥* 0 and takes values in 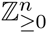 Thus, *X*(*t*) is a 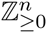-valued random variable, for each *t ≥* 0. Abusing notation, we also write *X*(*t*) to represent an outcome of this random variable on a realization of the process. The interpretation is:

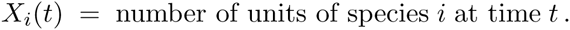

One is interested in computing the probability that, at time *t*, there are *k*_1_ units of species 1, *k*_2_ units of species 2, *k*_3_ units of species 3, and so forth:

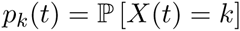

for each 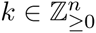. We call the vector *k* the *state* of the process at time *t*.

Arranging the collection of all the *p*_*k*_(*t*)’s into an infinite-dimensional vector, after an arbitrary order has been imposed on the integer lattice 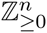, we have that 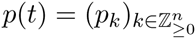 is the discrete probability density (also called the “probability mass function”) of *X*(*t*).

This note is concerned with the computation of moments, such as the expectation or mean (i.e, the average over all possible random outcomes) of the numbers of units of species at time *t*:

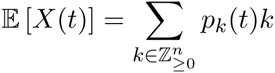

which is a column vector whose entries are the means

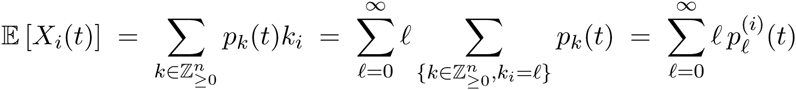

of the *X*_*i*_(*t*)’s, where the vector 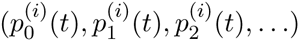 is the marginal density of *X*_*i*_(*t*). Also of interest, to understand variability, are the matrix of second moments at time *t*:

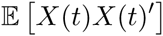

whose (*i, j*)th entry is E [*X*_*i*_(*t*)*X*_*j*_(*t*)] and the (co)variance matrix at time *t*:

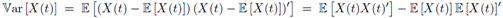

whose (*i, j*)th entry is the covariance of *X*_*i*_(*t*) and *X*_*j*_(*t*), 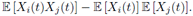

### 1.2 The Chemical Master Equation

A *Chemical Master Equation (CME)* (also known as a *Kolmogorov forward equation*) is a system of linear differential equations for the *p*_*k*_’s, of the following form. Suppose given *m* functions

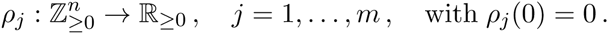

These are the *propensity functions* for the respective reactions *R*_*j*_. An intuitive interpretation is that *ρ*_*j*_(*k*)*dt* is the probability that reaction *R*_*j*_ takes place, in a short interval of length *dt*, provided that the state was *k* at the beginning of the interval. The CME is:

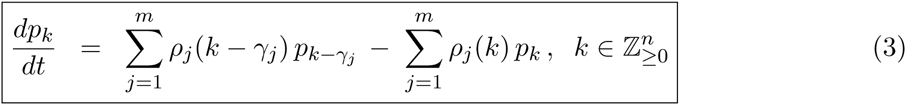

where, for notational simplicity, we omitted the time argument “*t*” from *p*, and where we make the convention that *ρ*_*j*_(*k - γ_j_*) = 0 unless *k ≥ γ_j_* (coordinatewise inequality). There is one equation for each 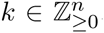, so this is an infinite system of linked equations. When discussing the CME, we will assume that an initial probability vector *p*(0) has been specified, and that there is a unique solution of (3) defined for all *t ≥* 0. A different CME results for each choice of propensity functions, a choice that is dictated by physical chemistry considerations. Here we will restrict attention to the most standard model, mass-action kinetics propensities.

We will also introduce the *n*-column vector:

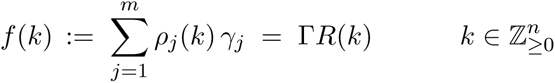

where *R*(*k*) = (*ρ*_1_(*k*), *…, ρ_m_*(*k*))^*1*^. One may interpret *f* (*k*)*dt* as the expected change of state during an interval (since *γ*_*j*_ quantifies the size of the jump if the reaction is *R*_*j*_). Thus, *f* (*k*) may be thought of as the rate of change of the state, if the state is *k*.

### 1.3 Propensity functions for mass-action kinetics

For each 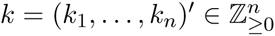, we let (recall that *a*_*j*_ denotes the vector (*a*_1*j*_, *…, a_nj_*)^*′*^):

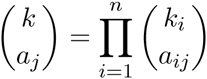

where 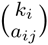 is the usual combinatorial number *k*_*i*_!/(*k*_*i*_ - *a*_*ij*_)!*a*_*ij*_!, which we define to be zero if *k*_*i*_ *< a*_*ij*_.

The most commonly used propensity functions, and the ones best-justified from elementary physical principles, are *ideal mass action kinetics* propensities, defined as follows:

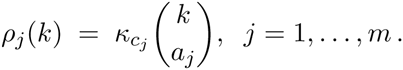

The *m* non-negative constants *κ*_1_, *…, κ_m_* are arbitrary, and they represent quantities related to the volume, shapes of the reactants, chemical and physical information, and temperature. Notice that *ρ*_*j*_(*k*) can be expanded into a polynomial in which each variable *k*_*i*_ has an exponent less or equal to *a*_*ij*_. In other words,

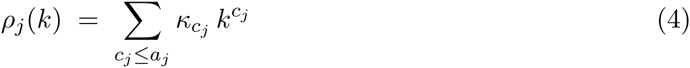

(“*≤*” is understood coordinatewise, and by definition 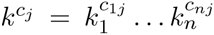 and *r*^0^ = 1 for all integers), for suitably redefined coefficients *κ*_*cj*_’s. Often one uses the simplification

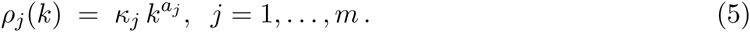

This simplification means that we approximate *x*(*x -* 1) *…*(*x - r* + 1) *≈ x^r^*, which introduces a small error if *r >* 1 and the integer *x* is very small. We provide results for both forms of propensity.

### 1.4 Moment equations

Suppose given a function 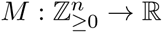 (to be taken as a monomial when computing moments). The definition of expectation gives:

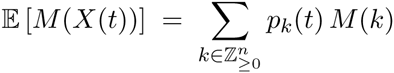

because ℙ [*X*(*t*) = *k*] = *p*_*k*_(*t*). We have:

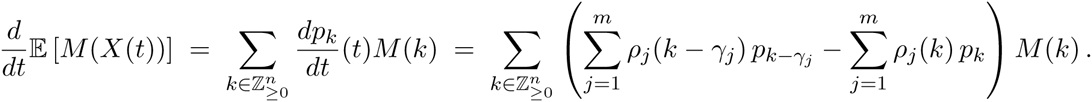

Note this equality, for each fixed *j*:

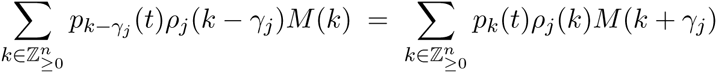

(by definition, *ρ*_*j*_(*k-γ_j_*) = 0 unless *k ≥ γ_j_*, so one may perform a change of variables 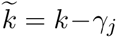). There results:

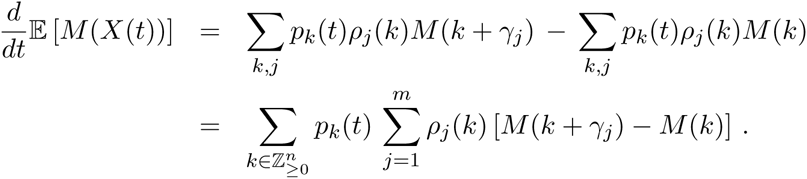

Let us define, for any 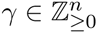, the new function Δ_*γ*_ *M* given by (Δ_*γ*_ *M*)(*k*) := *M* (*k* +*γ*) *- M* (*k*). With these notations,

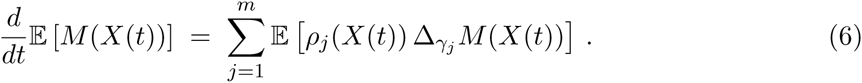

We next specialize to a monomial function:

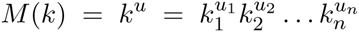

where 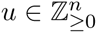. In this case,

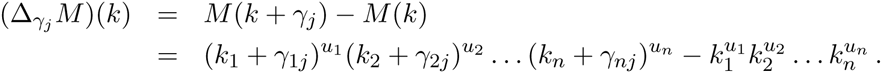

Expanding each binomial (*k*_*i*_ + *γ*_*ij*_)^*u*^*i* for which *γ*_*ij*_ ≠ 0, distributing, and canceling out the leading term, results in

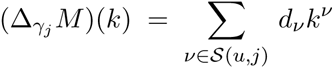

for appropriate coefficients *d*_*v*_, where

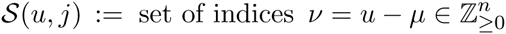

where 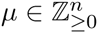 satisfies:

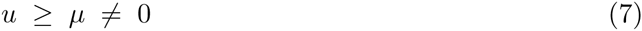

(inequalities “*≥*” in 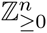 are understood coordinatewise), and

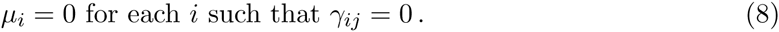

Thus, if 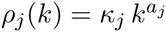 as in (5), then:

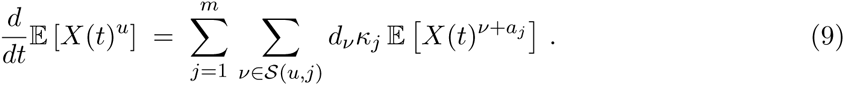

More generally, if

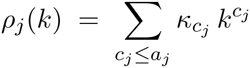

as in (4) then:

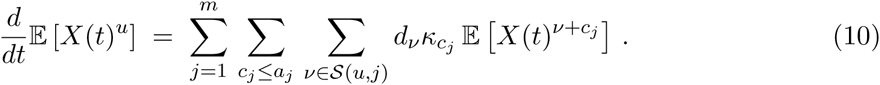

## 2 A set of transformations

For each multi-index 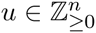, we define *R*^0^(*u*) = *{u}*,

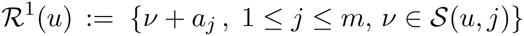

if using definition (5), or

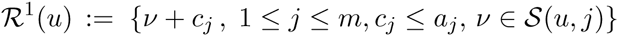

if using definition (4), and, more generally, for any *l ≥*1,

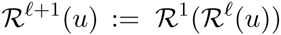

where, for any set *U*, *R*^*l*^(*U*) := ∪_*∈U*_ *R^l^*(*u*). Finally, we set

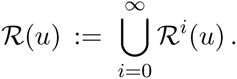

Each set *Rl*(*u*) is finite, but the cardinality #(*R*(*u*)) may be infinite. It is finite if and only if there is some *L ≥* 0 such that 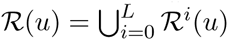, or equivalently 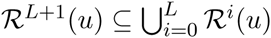.

Equation (9) (or (10)) says that the derivative of the *u*-th moment can be expressed as a linear combination of the moments in the set *R*^1^(*u*). The derivatives of these moments, in turn, can be expressed in terms of the moments in the set *R*^1^(*u*^*1*^), for each *u*^1^ *∈ R*^1^(*u*), i.e., in terms of moments in the set *R*^2^(*u*). Iterating, we have the following:

**Main Lemma.** Suppose that *N* := #(*R*(*u*)) *< ∞*, and write *R*(*u*) = *{u*_1_, *…, u_N_}*, with *u*_1_ = *u*. Then, there exists a matrix *A ∈* ℝ^*N ×N*^ such that, for any sample path *X*(*·*), and letting

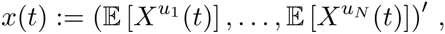

it holds that 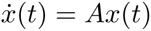 for all *t ≥* 0.

This motivates the following problem: *characterize those chemical reaction networks for which* #(*R*(*u*)) *< ∞ for all 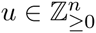.*

One trivial sufficient condition is that all reactions be or order 0 or 1, which means that *⊕a_j_ ∈ {*0, 1*}*. In that case, since *μ >* 0 in the definition of *S*(*u, j*), it follows that *⊕a_j_ ≤ ⊕μ* for every index *j*. Therefore, *⊕*(*ν* +*a*_*j*_) = *⊕u*+*⊕a_j_ -⊕μ ≤ ⊕u* for all *u*, and the same holds for *ν* +*c*_*j*_ if *c*_*j*_ *≤ a*_*j*_. In summary, the degree of all elements in *R*(*u*) is *≤ ⊕u*, so indeed #(*R*(*u*)) *< ∞*. A generalization to “weighted *L*^1^ norms” where one uses instead *⊕v* = *β*_1_*v*_1_ + *…*+ *β*_*n*_*v*_*n*_ for non-unity coefficients will be described next.

### 2.1 Lyapunov-like functions

**Definition.** A function: 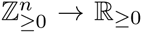 will be called a Lyapunov-like function with respect to a given chemical network if the following two properties hold:

1. for each *u, v*: *v ∈ R*^1^(*u*) *⇒ V* (*v*) *≤ V* (*u*) [nondecreasing property],
2. for each α *≥* 0: *V*_*a*_ := *{v | V* (*v*) *≤ a}* is finite [properness].

**Theorem.** For every chemical network, the following two statements are equivalent:

- There exists a Lyapunov-like function.
- #(*R*(*u*)) *< ∞* for all 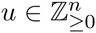.

**Proof.** Sufficiency is clear: pick any *u*, and let *α* := *V* (*u*); iterating on the nondecreasing property, *V* (*v*) *≤ α* for all *v ∈ R*(*u*), meaning that *R*(*u*) *⊆ V_a_*, and thus #(*R*(*u*)) *< ∞*.

To prove the converse, assume that #(*R*(*u*)) *< ∞* for all *u*. Define *V* (*u*) := max*w∈R*(*u*) *⊕w*. Since #(*R*(*u*)) *< ∞*, it follows that *V* (*u*) *< ∞*. As *u ∈ R*(*u*), it follows from the definition of *V* that *⊕u ≤ V* (*u*). Now pick any *u, v* so that *v ∈ R*^1^(*u*). Since *R*(*v*) *⊆ R*(*u*), it follows that *{⊕w, w ∈ R*(*v*)*} ⊆ {⊕w, w ∈ R*(*u*)*}*. Therefore *V* (*v*) *≤ V* (*u*) (nondecreasing property). Now pick any *α ≥* 0, which we may take without loss of generality to be a nonnegative integer, and any element *v ∈ V_a_*. Since *⊕v ≤ V* (*v*), it follows that *⊕ v ≤ a*. So *V*_*a*_ is a subset of the set of all nonnegative vectors *v* such that *⊕v ≤ a*, which has 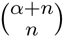 elements.

### 2.2 Linear *V*’s

The nonincrease requirement means, using the definition of *R*^1^(*u*), that

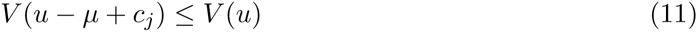

for all *c*_*j*_ *≤ a*_*j*_ (1 *≤ j ≤ m*) under definition (4) for propensities, or just for *c*_*j*_ = *a*_*j*_ if propensities have the simplified form (5), and every *μ* for which (7) and (8) hold, i.e., every *μ* for which *u ≥ μ ≠* 0 and *μ*_*i*_ = 0 for every *i* such that *γ*_*ij*_ = 0. Pick any reaction index *j* and for this index pick any species index *i* such that the species *S*_*i*_ changes, that is, *γ*_*ij*_ ≠ 0. Now pick *u* = *μ* = *e*_*i*_, the canonical unit vector with a “1” in the *i*th position (this choice of *μ* is allowed, since it is false that *γ*_*ij*_ = 0) and apply (11). that a necessary condition for decrease is that

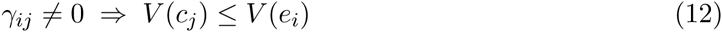

for all *c*_*j*_ *≤ a*_*j*_, or simply for *c*_*j*_ = *a*_*j*_ in the simplified case (5).

We now consider the special case of Lyapunov-like functions which can be extended to an additive map *V* : ℤ^*n*^ *→* ℝ. In this case, (12) equivalent to (11). Indeed, to see that (12) implies (11), pick any *u*, any reaction index *j*, and any *μ* such that (7) and (8) hold. Since *μ ≠* 0 and *μ ≤ u*, there is some species index *i* such that *γ*_*ij*_ ≠0 and *μ*_*i*_ ≠ 0, i.e., *μ ≥ e_i_*. Applying (12) with this choice of *i*:

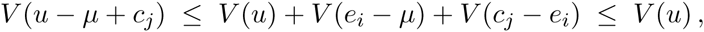

where we used that *V* (*μ - e_i_*) *≥* 0.

A map *V*: 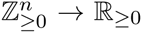 that extends to an additive function *V* : ℤ^*n*^ *→* ℝ is necessarily of the form

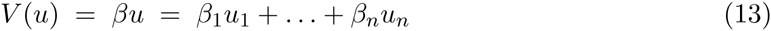

for some 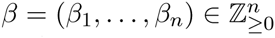 and it automatically satisfies the properness property provided that all *β*_*i*_ ≠ 0, which we assume from now on. Thus, #(*R*(*u*)) *< ∞* will be satisfied for all *u* if *V* has this form and satisfies (12). This condition can be made a little more explicit in the linear case. Let Δ_*j*_ := *{i | γ_ij_* =≠ 0*}*. Then a linear Lyapunov-like function amounts to picking a *β* such that

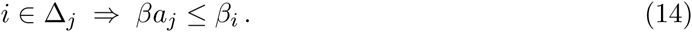

### 2.3 Special case: multi-layer feedforward networks

A general class for which there is a linear Lyapunov-like function, and hence #(*R*(*u*)) *< ∞* for all *u*, is that of multi-layer feedforward networks with linear reactions in the first layer. These are defined as follows. We find it convenient to separate degradations from more general reactions. So we will assume that there are reactions *R*_*j*_, *j ∈ {*1, 2, *…, m}*, which are partitioned into *p ≥* 1 layers: **R**_1_, *…,* **R**_*p*_. Species *S*_*i*_, *i ∈ {*1, 2, *…, n}*, are also partitioned into *p* layers **S**_1_, *…,* **S**_*p*_. In addition, we allow additional “pure degradation” reactions *D*_*j*_ : *S*_*i*_*j →* 0, *j ∈* {1, …, *d*} (so the total number of reactions is *m1* = *m* + *d*).

We assume that the reactions *R*_*j*_ that belong to the first layer **R**_1_ are all of order zero or one, i.e. they have *⊕a_j_ ∈ {*0, 1*}*. (This first layer might model several independent separate chemical subnetworks; we collect them all as one larger network.) More generally, for reactions at any given layer *π*, the only species that appear as reactants in nonlinear reactions are those in layers *< π* and the only ones that can change are those in layer *π*, that is:

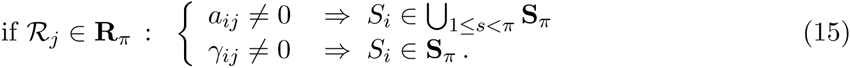

This means that except for order zero or one reactions, every reaction *R*_*j*_ at layer 1 *< π ≤ p* has the form:

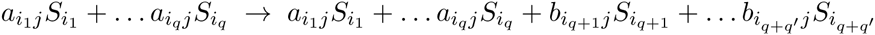

with *S*_*i*1_, …, *S*_*iq*_ in layers *< π* and *S*_*iq*+1_, …, *S*_*i*_*q*+*q1* in layer *π*. We claim that there is a linear Lyapunov-like function for any such network, Note that, for a degradation reaction *D*_*j*_ : *S*_*i*_*j →* 0, the entry *γ*_*ij*_ of the stoichiometry vector is nonzero (and equal to -1) only when *i* = *i*_*j*_, and for this index we have *a*_*ij*_ = 1. Thus condition (14) simply requires *β*_*i*_ ≤ *β*_*i*_ and hence is automatically satisfied no matter what is the choice of *β*. Thus we may ignore degradations and assume from now on that only the reactions *R*_*j*_ are present. We prove the claim by induction on the number of layers *p*. If *p* = 1, all reactions have order 0 or 1, so we can take *β*_*i*_ = 1 for all *i*. Arrange the species indices so that *S*_*r*+1_, …, *S*_*n*_ are the species in **S**_*p*_; these do not appear any reactions belonging to **R**_*π*_ for *π < p*. So layers **R**_*π*_ for *π < p* and species in **S**_*π*_ for *π < p* define a network with *p -* 1 layers, and we may assume by induction that a linear *V*_0_ has been defined for that network. This means that we have a vector of positive numbers *β*^0^ = (*β*_1_, *…, β_n-r_*) such that (14) holds for this subnetwork, which means, for any extension to a vector *β* = (*β*^0^, *\**) with *n* components (since the coefficients of *a*_*j*_ are zero for indices *r* + 1, *…, n*) that *βa_j_ ≤ β_i_* whenever *i ∈* Δ_*j*_, when *j*, *i* index reactions and species in the first *p -* 1 layers.

So all that is needed is to define the additional coefficients *β*_*i*_, *i ∈ {r* + 1, *…, n}*, such that the inequality *βa_j_ ≤ β_i_* holds for all pairs (*i, j*) such that (1) *R*_*j*_ *∈* **R**_*p*_ or *S*_*i*_ *∈* **S**_*p*_ and (2) *γ*_*ij*_ ≠ 0. We show that it suffices to pick all these *β*_*i*_ equal to a common value 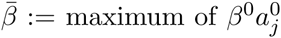 over all reactions *R*_*j*_ *∈* **R**_*p*_, where 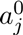 is the restriction of the vector *a*_*j*_ to its first *r* components. If *R*_*j*_ *∈* **R**_*p*_ and *S*_*i*_ /*∈* **S**_*p*_, the second condition in (15) (with *π* = *p*) says that (2) is not satisfied. Thus, we only need to consider *S*_*i*_ *∈* **S**_*p*_, i.e. *i ∈ {r* + 1, *…, n}*. Suppose first that *⊕a_j_ >* 1. The first condition in (15) (with *π* = *p*) insures that *a*_*ij*_ = 0 for all such *i*. Thus, 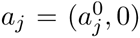 where the vector 0 has length *n - r*. it follows that 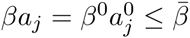. Next, suppose that *⊕a_j_ ≤* 1. If *⊕a_j_* = 0, then 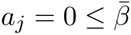. So assume *⊕a_j_* = 1 and pick the unique index *i*^*1*^ such that *a*_*i*_1*j* = 1. If *S*_*i*_1 *∈* **S**_*π*_, with *π < p*, then once again *a*_*ij*_ = 0 for all *i ∈ {r* + 1, *…, n}* and 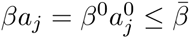. Finally, assume that *a*_*j*_ = *e*_*i*_ with *i ∈ {r* + 1, *…, n}*. Now 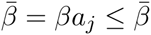 is trivially satisfied.

### 2.4 Examples

Let us start with the system shown in [1] to have moment closure:

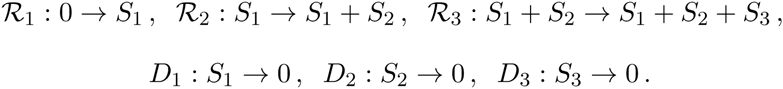

This is a three-layer system with one reaction in each layer, plus degradations. As we said, we may ignore degradations, so we consider:

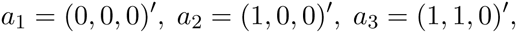

and we have Δ_1_ = *{*1*}*, Δ_2_ = *{*2*}*, Δ_3_ = *{*3*}*. We must find a positive vector *β* = (*β*_1_, *β*_2_, *β*_3_) such that *βa_i_ ≤ β_i_*, *i ∈ {*1, 2, 3*}*, i.e., *β*_1_ *≤ β*_2_ and *β*_1_ + *β*_2_ *≤ β*_3_. We may pick *β* = (1, 1, 2).

Here is a more complicated example involving several reversible first order reactions as well as some dimeric and trimeric reactions:

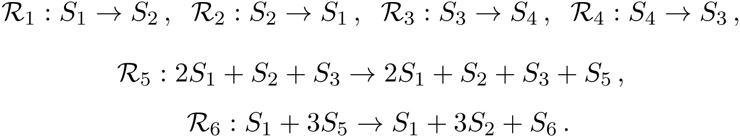

We have three layers: **R**_1_ = *{R*_1_, *R*_2_, *R*_3_, *R*_4_*}*, **R**_2_ = *{R*_5_*}*, **R**_3_ = *{R*_6_*}*, and **S**_1_ = *{S*_1_, *S*_2_, *S*_3_, *S*_4_*}*, **S**_2_ = *{S*_5_*}*, **S**_3_ = *{S*_6_*}*. Using *e*_*i*_ to denote canonical unit vectors:

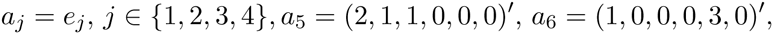

and

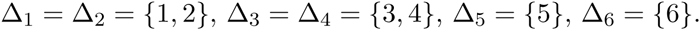

We must find a positive vector *β* such that *βa*_1_ *≤ β*_1_, *βa*_1_ *≤ β*_2_, *βa*_2_ *≤ β*_1_, *βa*_2_ *≤ β*_2_, *βa*_3_ *≤ β*_3_, *βa*_3_ *≤ β*_4_, *βa*_4_ *≤ β*_3_, *βa*_4_ *≤ β*_4_, *βa*_5_ *≤ β*_5_, *βa*_6_ *≤ β*_6_, i.e. so that *β*_1_ = *β*_2_, *β*_3_ = *β*_4_, 2*β*_1_ +*β*_2_ +*β*_3_ *≤ β*_5_, and *β*_1_ + 3*β*_5_ *≤ β*_6_. These constraints can be satisfied with *β*_1_ = *β*_2_ = *β*_3_ = *β*_4_ = 1, *β*_5_ = 4, *β*_6_ = 13.

### 2.5 Special case: conserved variables

In some applications, one is interested in computing the moments E [*X*(*t*)^*u*^] only for trajectories *X*(*t*) which remain in some specified subset 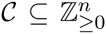. When this subset has the form

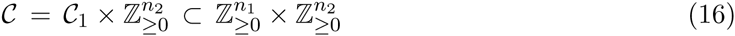

and the subset *C*_1_ is finite, the right-hand side of equation (10) (or (9)) can be simplified.

For example, suppose that the first two species *S*_1_ and *S*_2_ indicate the activity of a specified gene (inactive and active, respectively), with *S*_1_ and *S*_2_ reacting according to *S*_1_ *→ S*_2_, *S*_2_ *→ S*_1_, and no other reactions involve a change in *S*_1_ and *S*_2_. (This does not rule out reactions such as *S*_1_ *→ S*_1_ + *S*_3_ which would model transcription from the active conformation, since such a reaction does not change *S*_1_ nor *S*_2_.) It is the case that *X*_1_(*t*) + *X*_2_(*t*) remains constant in time, so *X*_1_(*t*) + *X*_2_(*t*) = *X*_1_(0) + *X*_2_(0) for all *t*. Moreover, given the biological motivation for these equations, we are only interested in the cases where (*X*_1_(0), *X*_2_(0)) = (1, 0) or = (0, 1). Thus, we have that *X*_1_(*t*) + *X*_2_(*t*) = 1 for all *t*. This restricts the components (*X*_1_(*t*), *X*_2_(*t*)) of *X*(*t*) to take values in the finite set *C*_1_ = *{*(1, 0), (0, 1)*}*, and hence all moments can be assumed to have the first two exponents equal to one:

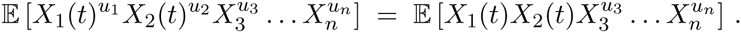

More generally, for any positive integers *r* and *L*, let *L*_*L,r*_ := *{*0, *…, L}^r^*. Then, for any finite subset 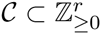, there is some integer *L* with the property:

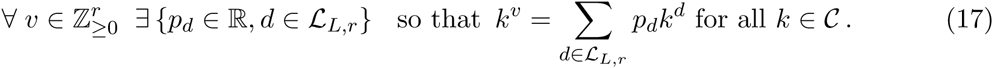

In other words, every monomial can be expressed as a linear combination of monomials with exponents *≤ L*.

To prove (17), observe that the set *F* of functions *C →* ℝ is a finite-dimensional vector space (canonically identified with ℝ^#(*C*)^, where #(*C*) is the cardinality of *C*). Introduce for each *i* the subspace *F*_*i,r*_ of *F* spanned by the monomial functions *k 1→ k^d^*, *d ∈ L_i,r_*. Since *F*_0,*r*_ *⊆ F*_1,*r*_ *⊆ F*_2,*r*_ *⊆ …*is a nondecreasing sequence of subspaces, there is some *L* such that *F*_*L*_1,*r* = *F*_*L,r*_ for all *L*^1^ *> L* (in fact, one may take *L* = #(*C*) - 1) and this proves the result.

A lower exponent may suffice for a proper subset of *{*0, *…, L}^r^*. For example, consider the set *C*_1_ = *{*(2, 0), (1, 1), (0, 2)*} ⊂ {*0, 1, 2*}*^2^. Then every monomial function on *C*_1_ is a linear combination of *f*_1_(*x, y*) = *x*, *f*_2_(*x, y*) = *xy*, and *f*_3_(*x, y*) = *y*. For example, *g*(*x, y*) = *x* 3 can be written as *g* = 4*f*_1_ - 3*f*_2_, as can be verified by plugging-in the three elements of *C*_1_. More generally, for *r* = 2 and two species satisfying *X*_1_ + *X*_2_ *≡ L*, we may write *X*_1_ = *L - X*_2_, and, using the inverse of the Vandermonde matrix of order *L* and evaluated at *{*0, *…, L}*, we may express 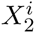 in terms of 1, *…*, 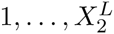, for any *i > L*.

Now given any set as in (16) with #(*C*_1_) *< ∞*, we may apply the above observation to *C*_1_, and this means that all moments 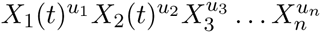 can be written as a linear combination of moments for which the first *n*_1_ exponents are *≤ L*. The remaining reactions could be a feedforward network, and now moments are all determined by a finite set of linear differential equations, so long as we only care about initial conditions in a finite invariant set. A simple example is as follows. We consider the following set of chemical reactions:

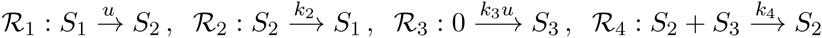

where we think of “*u*” as an external input. This is basically the incoherent feedforward loop considered in [2] to study adaptation and the fold-change detection property in stochastic systems. The only difference is that in that paper we used separate creation and degradation reactions 0 *→ S*_2_ *→* 0 (the first with rate *u*), but here, in order to impose a conservation law, we think of *S*_2_ as being an active form of a kinase (the input controlling the change to active form), which can be constitutively de-activated by a reverse reaction. The effect of *u* on *S*_3_ is incoherent, in the sense that *u* promotes formation of *S*_3_, as well as degradation, because the larger *u*, the larger the active concentration of *S*_2_, which degrades *S*_3_. Observe that we have

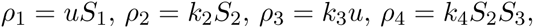

and

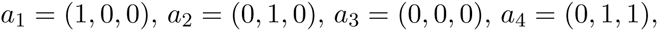

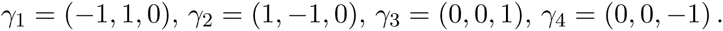

Note that *X*_1_(*t*) + *X*_2_(*t*) is constant along all solutions. Suppose that *X*_1_(0) + *X*_2_(0) = 2 (any number will work, but the formulas are more involved). Let us obtain a linear differential equation for the mean of *X*_3_(*t*). We introduce these notations:

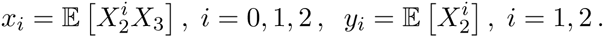

We are interested in *x*_0_(*t*). In general (omitting arguments *t*),

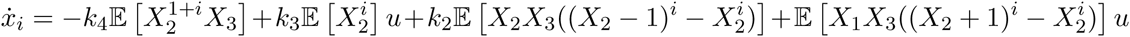

and

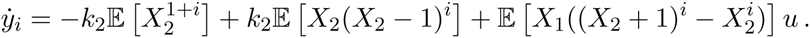

Using that *X*_1_ = 2 *- X*_2_ and 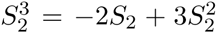 (which is valid whenever *S*_2_ *∈ {*0, 1, 2*}*), we conclude that:

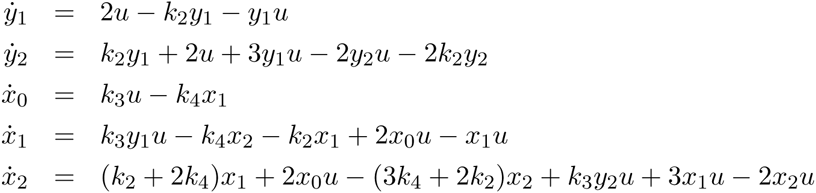

which may be written in the “bilinear” form 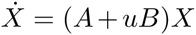, where *X* = (*y*_0_, *y*_1_, *y*_2_, *x*_0_, *x*_1_, *x*_2_)^*1*^with the convention that *y*_0_ ≡ 1 and appropriate 5 × 5 matrices *A, B*. Similar (but larger) systems may be written for the second and larger moments of *X*_3_ as well as mixed moments.

More abstractly, given any finite continuous-time Markov chain with *n*_1_ states *q*_*i*_ and transition rates *λ*_*ij*_, we may introduce *n*_1_ species *S*_*i*_ and reactions *S*_*i*_ → *S*_*j*_ with rate *λ*_*ij*_. The stoichiometric matrix consists of columns with exactly one entry equal to 1 and one entry equal to -1, so the sum *X*_1_(*t*) + *…*+ *X*_*n*1_ (*t*) is conserved (see e.g. Section 4.8 in [3]). Thus, starting from an initial condition with *X*_1_(0) + *…*+ *X*_*n*1_ (0) = 1 we have that at all times we have precisely one *X*_*i*_(*t*) = 1. This provides an embedding of the Markov Chain: state is *q*_*i*_ at time *t* if *X*_*i*_(*t*) = 1. This construction is of interest when reaction parameters *κ*_*i*_ in a network are described by functions of finite Markov chains (Hidden Markov Models) and the network is of a feedforward type, to conclude that finite-dimensional ODE’s exist for moments.

